# Reproductive barriers and genomic hotspots of adaptation during allopatric species divergence

**DOI:** 10.1101/2022.03.11.483945

**Authors:** Riddhi Deshmukh, Saurav Baral, Muktai Kuwalekar, Athulya Girish Kizhakke, Krushnamegh Kunte

## Abstract

Theory predicts that in allopatric populations, genomic divergence and reproductive barriers will be driven largely by random genetic drift, and thereby evolve slowly in large populations. However, local adaptation and divergence under selection may also play important roles, which remain poorly characterised. Here we address three key questions in young allopatric species: (a) How widespread are genomic signatures of adaptive divergence?, (b) What is the functional space along which young sister species show divergence at the genomic level?, and (c) How quickly might prezygotic and postzygotic reproductive barriers evolve? Analysis of 82 re-sequenced genomes of the Oriental *Papilio polytes* species group revealed surprisingly widespread hotspots of intense selection and selective sweeps at hundreds of genes unique to each species, and spanning all chromosomes, rather than divergence only in a few genomic islands. These genes perform diverse ecologically important adaptive functions such as wing development, colour patterning, courtship behaviour, mimicry, pheromone synthesis and olfaction, and host plant use and digestion of secondary metabolites, that could contribute to local adaptation and subsequent reproductive isolation. Divergence at such functional genes appeared to have reproductive consequences: behavioural and hybridisation experiments revealed strong assortative mate preference (prezygotic barriers) as well as postzygotic barriers to hybridisation in timespans as short as 1.27 my, indicating that speciation was already complete, rather than incipient. Our study thus demonstrates an underappreciated role of intense selection and potential local adaptation in creating genome-wide hotspots of rapid molecular evolution and divergence, during differentiation and speciation in young allopatric species.

## Introduction

The interaction between speciation and diversification has shaped the remarkable diversity of lifeforms. The causes and consequences of selective agents and processes that drive this diversification are critical in structuring biological diversity. Research in this area was historically disconnected to a degree. Traits within a species were studied largely through the lens of selection and local adaptation^1–4^. On the other hand, speciation was studied largely through the lens of geographic isolation, in which drift is expected to play a prominent role in the evolution of reproductive isolation^5^. This was unfortunate because adaptative divergence and ecological, physiological and genetic bases of reproductive isolation were very much at the core of early thinking about speciation^1,2,6,7^. These two views began to coalesce with the renewed interest in the process of adaptive divergence and speciation, and the availability of genetic and genomic data^7,8^. The recent shift towards understanding the biological aspects, rather than geographic aspects, of divergence and speciation has generated a robust understanding of how ecological pressures create disruptive natural and sexual selection. These disruptive selections fuel adaptive divergence along morphological, behavioural, sensory and physiological axes across the population-to-species continuum. Some of the prominent examples of adaptive population divergence along this continuum are: (a) sympatric host race formation in pea aphids^9^ and *Timema* stick insects^10^, and in divergent male courtship signals and female preference in bushcrickets (*Ephippiger diurnus*)^10^ and frogs (*Engystomops petersi*)^11^; (b) at early stages of reproductive isolation and speciation, polygenic basis of reproductive isolation involving hybrid male sterility and genomic regions with strong differentiation in *Oryctolagus cuniculus* rabbits^12^; (c) incipient ecological speciation primarily due to local adaptation to water availability influencing several morphological traits in subspecies of *Boechera stricta*, a perennial mustard^13^; and at the other end, (d) clear reproductive isolation in well-established sister species, e.g., assortative mating by wing colouration among *Heliconius cydno* and *H. pachinus*, governed by a single locus that controls both wing colour pattern and mate preference^14^. Similarly, pollinators mediate prezygotic isolation in *Mimulus lewisii* and *M. cardinalis*, the underlying flower colour being controlled by a major quantitative trait locus^15^. Work in this area has so far focused largely on the tempo and mode of graded divergence, gene flow, and genetics of ecological adaptation and reproductive isolation along the population-to-species continuum^16,17^. However, relatively little is known, and from only a few intensively studied organisms^18,19^, about broad genome-wide signatures of genetic incompatibilities, adaptive divergence, and the roles of selection and drift in shaping this genomic divergence among species.

There is value in understanding the interaction between ecological pressures and geographic isolation, and the population genetic signatures they may leave on adaptation and genomic divergence during the process of speciation. In allopatric populations, one might expect neutral processes such as genetic drift to be prominent in driving genomic divergence among populations^5,20^, resulting in slower evolution of reproductive isolation^21–23^. On the other hand, genetic/genomic divergence and reproductive isolation may be accelerated in sympatric populations evolving under selection and adaptive divergence^21– 23^. Thus, although the focus of investigations should rightfully be on the nature and cause of selection and divergence, the geographic setting offers good natural experiments that may be used to ask interesting questions about ecological and genomic divergence. For example, to what extent is genomic divergence influenced by drift, local adaptation, or ecological divergence and reproductive isolation *per se* that might maintain co-adapted phenotypic and gene complexes.

Here we focus on the tempo and genomic consequences of divergence between young species. Divergence predominantly under drift versus selection may produce different genomic signatures. Under the drift scenario, there may be weak selection for divergence *per se*. Therefore, populations may diverge slowly, producing few hotspots of adaptive divergence and little reproductive isolation, and certainly not prominent signatures of evolution under intense selection. On the other hand, populations diversifying under local selection and adaptive divergence may produce distinct genomic signatures of positive and directional selection or, in extreme cases, hard selective sweeps, for traits that are under selection. Under the local selection and adaptive divergence scenario, one would still expect the majority of the genome to evolve under drift according to the neutral theory of molecular evolution^24^. Nonetheless, the genomic signatures of divergence under selection would still be prominent under this scenario and not under the drift scenario. Indeed, certain ecological adaptations appear to be primed for rapid divergence and local adaptation in the process of population differentiation and speciation, perhaps because they often experience intense selection and have malleable genetic basis that responds to selection rapidly. Known examples include colour polymorphisms, mimicry, armature and other traits used in predator avoidance^25– 28^, diet specialisation^9,29,30^, and sexually selected adaptations such as courtship behaviours, sexual weapons and ornaments^31–33^. Therefore, we address two key questions: (a) How widespread are the genomic signatures of adaptative divergence?, and (b) What is the functional space (i.e., range of functions performed by genes involved in ecological adaptations) along which sister species show prominent genomic divergence?

We explore this framework by analysing genomes of the *Papilio polytes* species group. The *Papilio polytes* group has recently emerged as a model system in evolutionary and developmental genetics because of its iconic mimetic polymorphism^34–37^. *Papilio* are edible prey but females of many species have evolved into Batesian mimics of the aposematic *Pachliopta* butterflies to avoid predators^38,39^. Throughout its wide range in the Indo-Australian Region, males of *P. polytes* in all populations have a single non-mimetic, white-banded colour form but their females have four forms: *f. cyrus* is male-like and non-mimetic, *f. polytes* and *f. theseus* mimic the white-spotted and black forms of *Pachliopta aristolochiae*, and *f. romulus* mimics *Pachliopta hector* (Fig. 1). Forms *cyrus* and *polytes* co-occur across large part of the range of these species, while *f. romulus* is restricted to Peninsular India and Sri Lanka, and *f. theseus* is endemic to the islands of SE Asia^40^. Because of this single male and multiple female forms, the species group is popularly known as Mormon swallowtails. Historically, the *Papilio polytes* group was considered to have only three species—*P. polytes, P. phestus* and *P. ambrax* (Fig. 1a). Of these, *P. polytes* was believed to be the most widespread, occurring from Sri Lanka and India to the Lesser Sunda Islands, Sulawesi, the Philippines and southern Japanese islands in the east, splintered into at least 20 subspecies structured in the mainland-island mosaic and exhibiting small variation in wing colouration and the presence/absence of tails^40–42^. However, there has been understated evidence indicating some reproductive incompatibility between a few subspecies. For instance, Clarke and Sheppard had different degrees of success in crossing several putative subspecies of *P. polytes* to identify the genetic bases of mimetic wing polymorphism^40^. Conflicting phylogenetic evidence has also recently emerged. Reconstruction of the evolutionary history of mimetic polymorphism in this group with genome-wide markers showed *P. polytes* to be either a polyphyletic or a paraphyletic species because *alphenor*, traditionally considered to be the Philippine subspecies of *P. polytes*, was sister to *P. phestus* and *P. ambrax* outside of *P. polytes*^43^ (Fig. 1b). Further, our preliminary subspecies-level mito-nuclear phylogeny of the *polytes* species group suggested that *P. polytes* is not one but a complex of three geographically well-separated (allopatric) species—*P. polytes, P. javanus* and *P. alphenor*—each with multiple island subspecies included in them (Fig. 1c–d). The time calibrated phylogeny and node age estimates suggested the group to have evolved approx. 4.69 mya with the split between the youngest *P. polytes* and *P. javanus* species pair being 1.27 mya. Thus, the systematics of this complex polytypic group is still unresolved, requiring a more thorough genome-level analysis of selection, divergence and differentiation, along with an understanding of reproductive barriers.

**Fig. 1:**
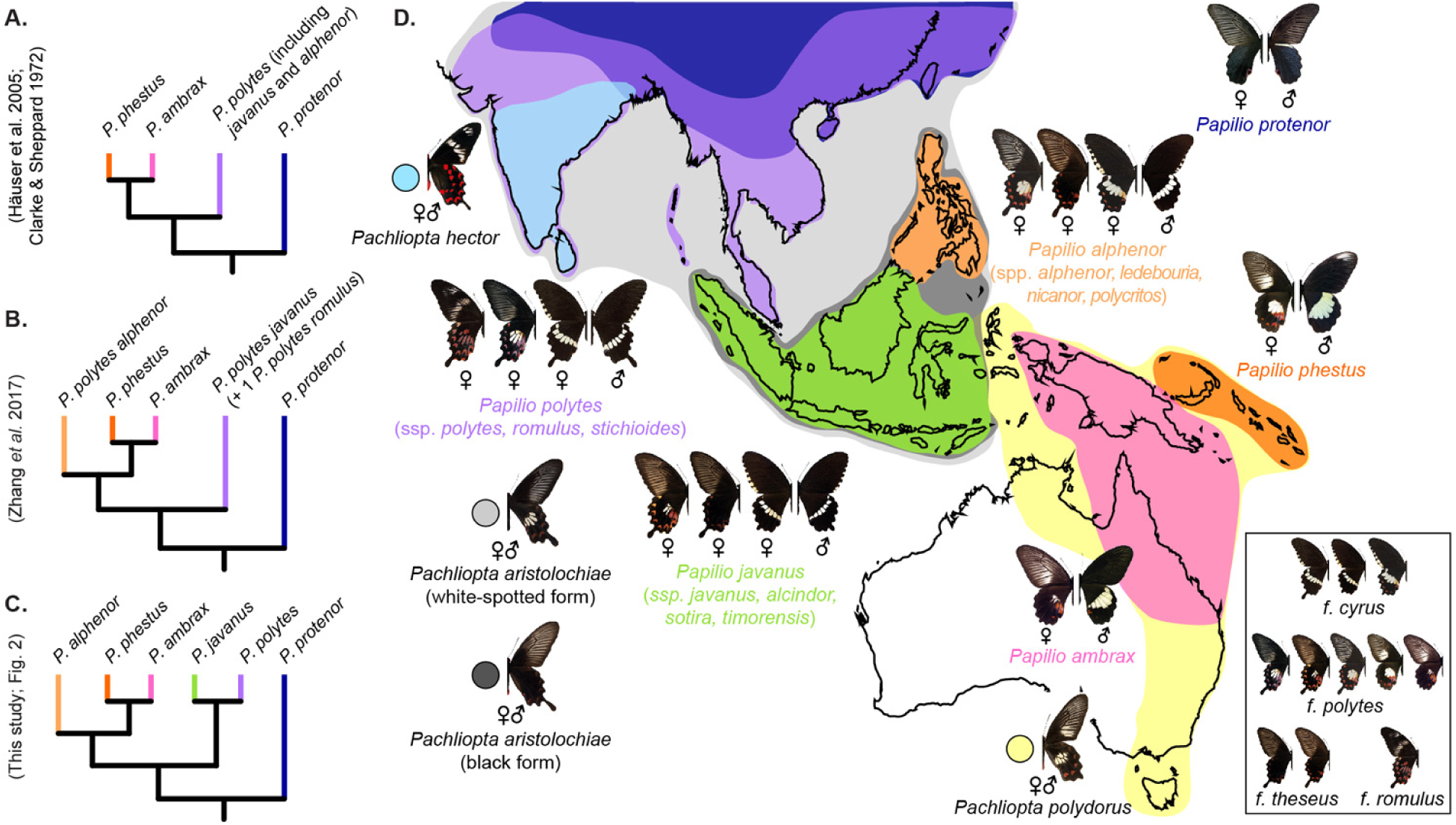
Putative species relationships, geographical distribution and various Batesian mimetic forms in the *Papilio polytes* species group. A–C. Different interpretations of species relationships in the *polytes* species group. **D**. Geographical ranges of each species, their sexual dimorphism, female polymorphisms, and Batesian mimetic relationships. Names of all aposematic models are in black, whereas names of all the *polytes* group species are colour coded with their geographic ranges as per latest taxonomic proposals (this work). Sexual dimorphism, polymorphism and mimicry phenotypes are also illustrated for each taxon.

Here, we first characterise species in the *P. polytes* species group with phylogenetic, genomic and behavioural analyses. We then use this characterisation to address the following questions: (a) Is *P. polytes* a polyphyletic, paraphyletic or monophyletic species? (b) If it is indeed a complex of three well-defined allopatric species as suggested recently, do their genomes show prominent signatures of divergence under selection or drift? (c) Are signatures of selective divergence concentrated in a few genomic islands or is divergence more widespread at the genomic level?, and (d) What is the functional space of genes that show signatures of selective divergence? Are these shared between species? In addition to illuminating aspects of selection, divergence and reproductive isolation among these allopatric species, we present the implications of our results for systematics of polytypic species in the Indo-Australian Region.

## Results

### *Papilio polytes* is a cluster of three well-supported, monophyletic, geographically structured species. *Phylogenetic structure*

We generated Bayesian phylogenies of the *polytes* species group, including multiple subspecies, with four marker datasets (Fig. 2). A species tree reconstructed using only the mitochondrial markers was unresolved (Fig. 2d), showing that barcodes and other mitochondrial markers alone are not adequate to resolve relationships in this species group. However, the three trees reconstructed using the nuclear and mito-nuclear datasets produced an identical, well-supported topology showing the following species relationships:

**Fig. 2:**
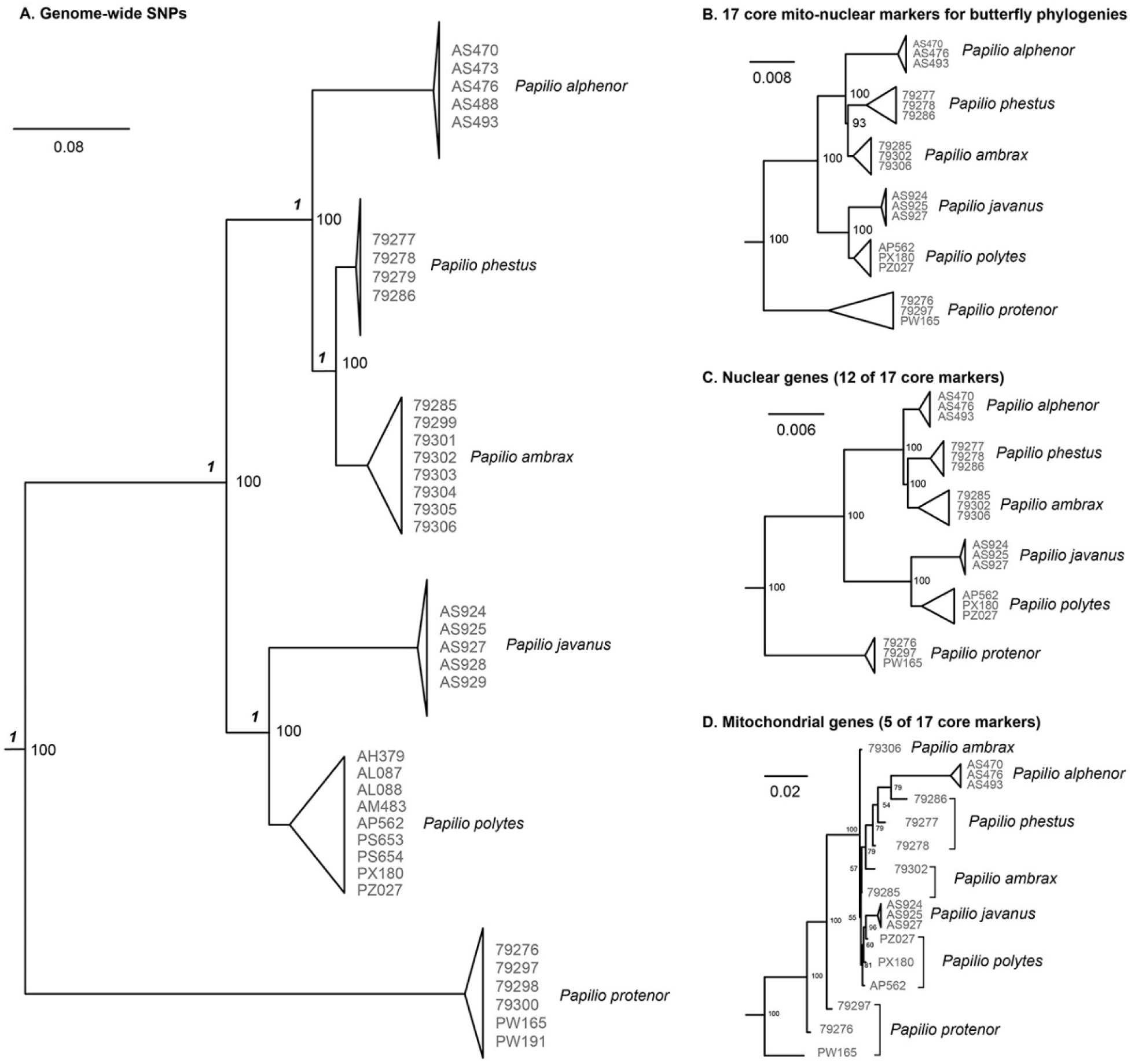
Species phylogenies of the *polytes* species group using different sets of genetic markers. Numbers adjacent to collapsed branches represent specimen codes or genome accession numbers (sample details in Table S1). Support values for all splits are shown at the nodes. Italicised numbers in (A) indicate mPTP support for species divergence events. The use of only mitochondrial markers resulted in an unresolved tree topology (D).

> (*protenor*,((*alphenor*,(*phestus, ambrax*)),(*polytes, javanus*))) (Fig. 2A–C).

Thus, *P. polytes* as traditionally viewed came out as a polyphyletic species. The three distinct, strongly supported monophyletic groupings that resulted, which do not even show sister species relationships with respect to *alphenor*, can be treated as three distinct species, as *P. polytes, P. javanus*, and *P. alphenor*. The tree topology, node age estimates, mainland-island distributions, and allopatry of these three monophyletic groups, all indicate that the three species have diverged in allopatry over the past 4.69 million years, i.e., since early Pliocene.

#### Species delimitation

We performed species delimitation analysis with SNAPP using the 30,760 genome-wide SNP dataset (dataset 1). We tested five models with different combinations of species partitions (Table 1) against a null model assuming that *P. polytes, P. javanus* and *P. alphenor* are distinct species (based on Figs. 1C and 2A–C). We derived the alternative models based on previous taxonomic understanding of species relationships in the *P. polytes* species group (models B–C), geographical landscape over which these taxa occur (models E–F), and results of the population structure analysis from the next section (model D). None of the alternative models obtained sufficient support: models B–F were rejected (Bayes factor<0) in favour of the null model with the highest MLE rank (Table 1). We ascertained the most supported species partitions using mPTP (multi-rate Poisson Tree Processes model) analysis, which models speciation rate using sequence substitutions. This analysis also strongly supported the topology in Fig. 2A–C, that *P. polytes, P. javanus* and *P. alphenor* are three distinct, monophyletic species (mPTP support values in italics in Fig. 2A).

**Table 1:** Comparison of alternative species models with SNAPP. Marginal Likelihood Estimates (MLE) and Bayes factors for different models of species delimitation are shown below. The null model for calculation of Bayes factor was A, where *P. polytes, P. javanus* and *P. alphenor* were considered different species with the species topology as shown in Fig. 1.

| Model | No. of species | MLE | MLE rank | Bayes factor |
| --- | --- | --- | --- | --- |
| A (*polytes* species group has 6 species; Fig. 1C, 2A–C) | 6 | -119050 | 1 | - |
| B (*polytes* and *javanus* are one species; Fig. 1B) | 5 | -137543 | 5 | -36986.268 |
| C (*polytes*, *javanus* and *alphenor* are one species; Fig. 1A) | 4 | -195363 | 6 | -152626.3 |
| D (*ambrax* and *phestus* are one species; Fig. 3) | 5 | -127348 | 3 | -16596.529 |
| E (*polytes* split into two species, one occurring in South Asia (India) and the other in Indo-China (Thailand)) | 7 | -121271 | 2 | -4441.1624 |
| F (Thailand *polytes* + *javanus*) | 6 | -137311 | 4 | -36522.321 |

#### Population structure

We used ADMIXTURE to determine population structure in the *polytes* species group using biallelic genome-wide SNPs. The most supported result with the smallest cross-validation error split the group into six clusters (K=6; Fig. S2B). *Papilio polytes, P. javanus* and *P. alphenor* formed separate population clusters, with *P. polytes* being split into two distinct clusters over its geographic range. Such a clear population structure in *P. polytes* appears to be a result of considerable genomic variation in its large population size over a very broad geographic range.

ADMIXTURE analysis treated *phestus* and *ambrax* together as a single population cluster at K=5– 7. However, the SNAPP model that assumed *phestus* and *ambrax* to be a single species (model D) was not well supported (Table 1), and species delimitation analysis using mPTP also showed them to be strongly supported taxa at a species level, so we treat them as such (Fig. 2A–C).

#### Genome-wide divergence

These species show considerable percentage divergence between the genome-wide variable sites (30,760 SNPs from dataset 1; range: 3.6–17% divergence; Figs. 2A, S1). We further estimated pairwise genomic divergence (Fst) in 10kb windows using the full dataset of 27 million genome-wide SNPs across the five mimetic sister species (Fig. S3). Genome-wide Fst was considerably high, ranging from 0.4 to 0.8, revealing high divergence among all the species (Fst values: *P. polytes–P. javanus*: 0.43; *P. polytes–P. alphenor*: 0.64; *P. javanus–P. alphenor*: 0.82; *P. phestus–P. ambrax*: 0.59; Fig. S3). It is evident from the Fst values that the level of genome-wide divergence was not correlated with the phylogenetic or geographic distance between species, perhaps indicating that the divergence might not have been driven purely by neutral processes. Although effective population size and gene flow may influence the pattern of genetic divergence observed, the three species have fairly large and widespread populations across their geographical ranges^43^, and they occur in allopatry with no known hybrid zones (Fig. 1D).

### *Papilio polytes, P. javanus* and *P. alphenor* are reproductively isolated by prezygotic and postzygotic barriers

We tested mate preference (prezygotic barriers) and post-mating reproductive success (largely postzygotic barriers) using naturally occurring and hand-paired intraspecific and interspecific matings in the three taxa (*polytes, javanus* and *alphenor*) that are still widely treated as subspecies of the same species. In the first experiment in a mixed population of freshly eclosed individuals (see Methods), all three species mated assortatively (Pearson’s Chi-squared test, χ^2^=100.44, df=4, p<0.0001; Fig. 3A, Tables S3, S5) even when they had no prior experience of their own species over other species, showing that the assortative mate preference was instinctive. In the second set of experiments, we tested reproductive success by setting up intraspecific and interspecific hand-paired matings. This experiment removed the behavioural choice of mates but was able to reveal post-mating selection, reproductive compatibilities and hybrid fitness. We compared several measures of reproductive success: (1) the duration of mating, (2) no. of eggs laid, (3) no. of eggs that hatched successfully, and (4) no. of adults that eclosed successfully. We compared the reproductive success in hand-paired intraspecific matings with those of naturally occurring intraspecific matings. Naturally paired and hand-paired intraspecific matings showed similar trends in all the traits measured (Tukey’s test with correction for multiple comparisons, p>0.08 for all pair-wise comparisons; Fig. 3B–C, Tables S2, S4–6), showing that hand-pairings were successful. Likewise, mating duration as well as the number of eggs laid between intraspecific and interspecific matings were similar (Tukey’s test with correction for multiple comparisons for all pair-wise tests: mating duration: p>0.8, except *polytes-alphenor* hand-paired mating where p=0.04; no. of eggs laid: p>0.1), further showing that hand-pairings resulted in normal mating and oviposition behaviours. This result also suggests that the prezygotic barriers between these species do not appear to affect the maturation and oviposition of eggs. However, the hatching and eclosion success in hand-paired interspecific matings was very low (Tukey’s test with correction for multiple comparisons, p<0.001 for all pair-wise tests; Fig. 4D–E, Tables S2, S4–6), showing that the three species have genetic/developmental incompatibilities that contribute to postzygotic barriers to hybridisation.

**Fig. 3:**
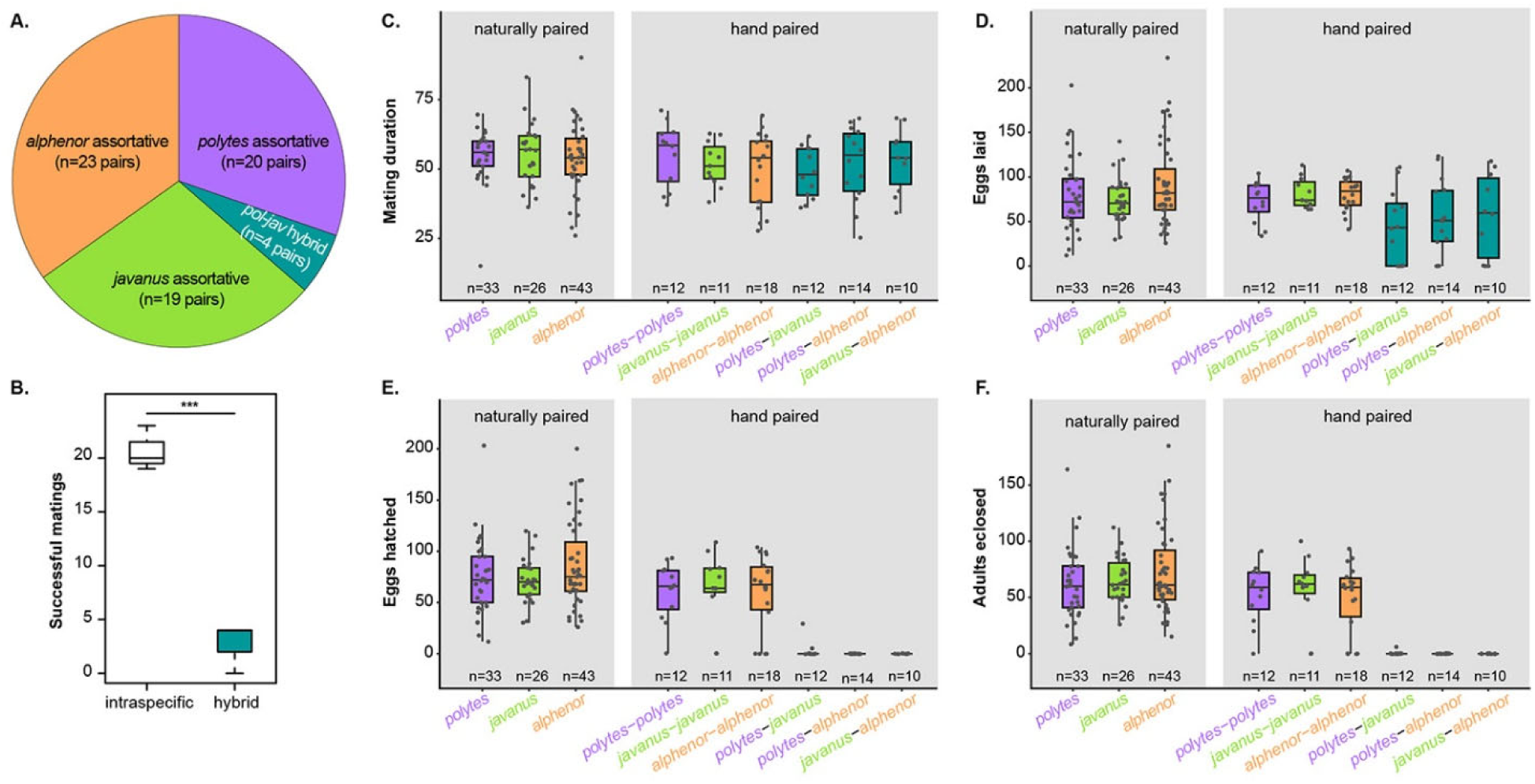
Reproductive success and isolation among *P. polytes, P. javanus* and *P. alphenor*. A–B. Test of prezygotic barriers to reproduction. Assortative mating in a mixed population containing equal numbers of male and female of *P. polytes, P. javanus* and *P. alphenor*. **C–F**. Tests of postzygotic barriers to reproduction. Mating duration (C), fecundity (D), hatching success (E), and eclosion success (F) across the three species and their hybrids in naturally-paired and hand-paired matings. p<0.001, Tukey’s test with correction for multiple comparisons, for all pair-wise tests between intraspecific and interspecific matings in (E) and (F). Details of the statistical tests are provided in Table S6. Intraspecific matings are colour coded, interspecific matings are dark green.

**Fig. 4:**
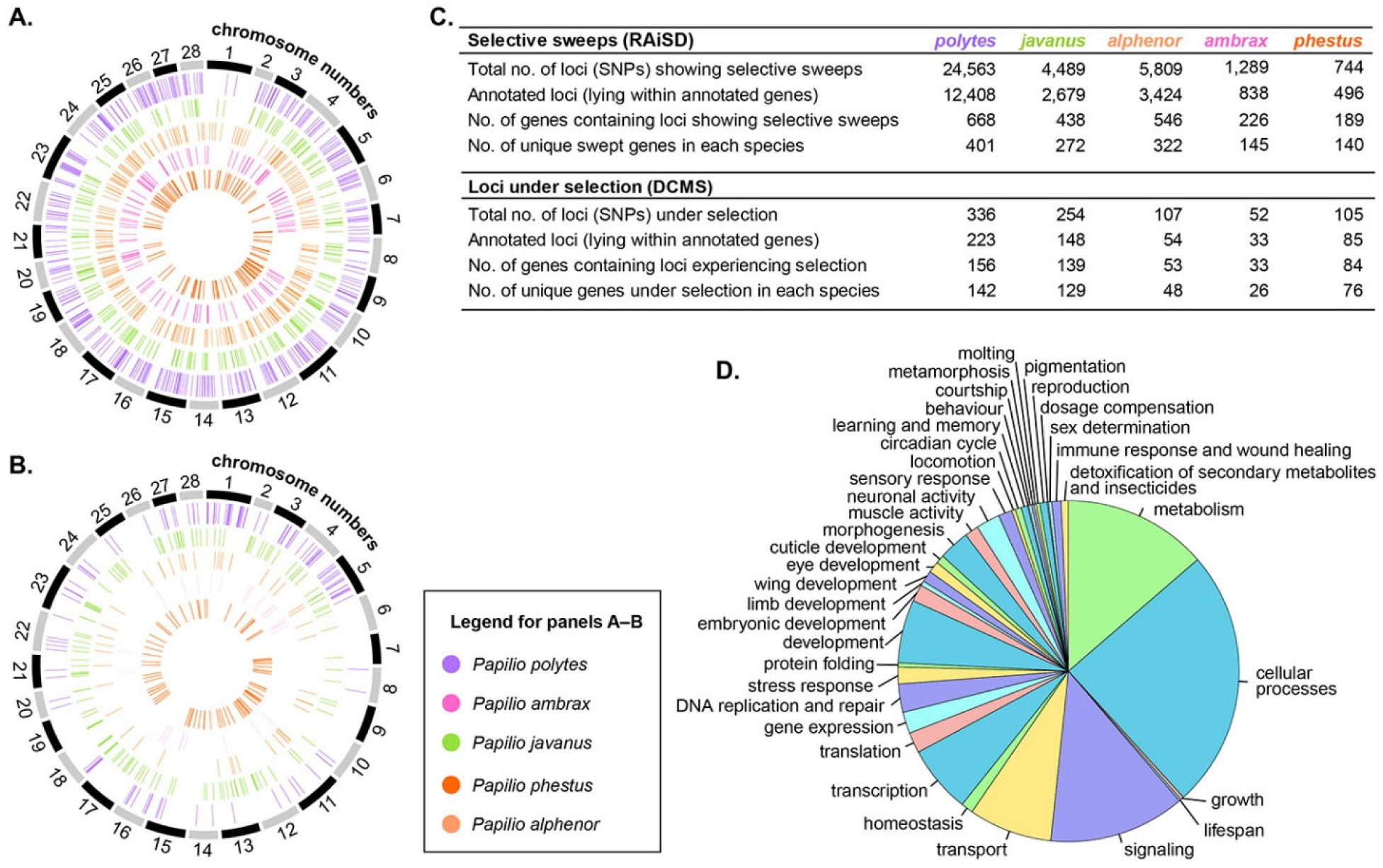
Loci under selection during genomic divergence of species in the *P. polytes* species group. A. Annotated genes showing selective sweeps in the five species, identified using RAiSD. The alternately coloured outer bands indicate chromosomes as defined in the reference *Bombyx mori* genome, whereas colour-coded lines inside represent genes under selection unique to each species. **B**. Annotated genes showing signatures of selection, identified using DCMS. Colour coding as in panel A. **C**. Summary of the number of loci (SNPs and the genes they lie in) experiencing intense selection in the *P. polytes* species group. **D**. Range of functions performed by genes containing sites under selection in the five *Papilio* species. Details of gene functions are provided in Table S10.

### All *P. polytes* group species show widespread genomic signatures of strong selection and divergence

We predicted that if the *P. polytes* group species have largely drifted apart in allopatry, their genomes would likely show little signatures of divergence under selection. On the other hand, if their divergence has occurred under some local selection, then we may observe genomic signatures of divergence under selection and adaptation. To test whether the *P. polytes* group species show prominent signatures of selection, we characterised the genomic regions that have experienced intense selection since the species started evolving independently after initial geographic isolation. We used Raised Accuracy in Sweep Detection (RAiSD) to identify selective sweeps, and a composite measure of selection (de-correlated composite of multiple signals (DCMS), including H1, H12, Fst, Tajima’s D and nucleotide diversity), to identify regions showing signatures of selection across the genomes. We particularly chose signatures of selective sweeps as a clear indication of intense selection and its genetic manifestation, whereas the composite measure of selection may indicate a variable intensity of selection, from weak to strong. Together, these two approaches revealed that: (a) hundreds of genes have experienced selective sweeps or diversifying selection in each species, (b) a large proportion of genes was under selection in individual species of the group, and (c) few genes were under selection in more than two or three species (Fig. 4A; Tables S7–S8). Moreover, the sets of genes under selection as identified by RAiSD (selective sweeps) and DCMS (selection) were completely non-overlapping (Tables S7–S8). Only the mimicry locus (*doublesex*)^44,45^ showed signatures of selective sweeps in all five mimetic species, whereas DCMS did not identify any gene under selection in all species. The genes experiencing selective sweeps/selection were not concentrated in a few genomic islands but spread widely in numerous hotspots across all the annotated chromosomes in the genome (Fig. 4A–C). Such a broad signature of genome-wide divergence under selection is particularly striking in light of the relatively recent splits between these species: the oldest split in this group was approx. 4.69 mya, between (*polytes, javanus*) and (*alphenor*,(*ambrax, phestus*)), and the most recent splits were only approx. 1.27 mya between (*polytes, javanus*) and approx. 1.95 mya between (*ambrax, phestus*). Moreover, the fact that the genome of each of these young allopatric species shows such widespread signatures of selective sweeps itself is startling.

### *Papilio polytes* group species have diverged under selection on hundreds of genes involved in local adaptation, sexual selection, and housekeeping

Species of the *P. polytes* group show two striking patterns: (a) all the species have non-overlapping geographic distributions over different mainland/island groups (Fig. 1D), and (b) a large number of genes are under selection in individual species (Fig. 4C, Table S9). These observations together imply that, contrary to the expectation of slow, largely neutral divergence under drift in allopatric populations, these allopatric species have diversified under considerable selection as shown by the widespread hotspots of genomic divergence. What is the functional space of genes that show divergence under selection in these species? We mapped all the SNPs experiencing selection to annotated genes with known functions to characterise the functional space that these genes occupy. These genes belong to the following major functional categories: (a) local adaptation, such as host plant use, digestion of secondary metabolites, and insecticide resistance, (b) mate recognition and reproductive isolation, such as wing patterning, eye development, pheromone production, odorant receptors, and courtship behaviour, (c) embryonic and organismal development, and gene expression, and (d) cellular housekeeping genes involved in metabolism, transport and signaling (Fig. 4D, Tables S7, S8, S10). Of these, the first two functional categories have clear implications for ecological and sexual selection, and local adaptation and divergence. Indeed, the pre- and postzygotic barriers to hybridisation that we uncovered above, such as assortative mating and low reproductive success in hand-paired interspecific crosses, may have direct links to functional isolation along sensory axes, facilitated by mate choice, and by physiological and developmental breakdown in hybrid zygotes. A closer look at the biology of these species and additional hybridisation experiments may shed further light on this.

Of particular interest is the gene *doublesex*, which shows signatures of selective sweeps in all five mimetic species of the group but not in the basal non-mimetic *P. protenor*. It is important not only in mimetic polymorphism and wing patterning in the *polytes* species group but, as a key developmental regulator of sexual dimorphisms, it may be involved in other sex-specific adaptations in these species. The last two functional categories are important for a swath of critical functions at an organismal level but it is unclear how selection on these genes might contribute to local adaptation and divergence. This needs to be explored further.

## Discussion

Isolation, divergence and speciation in the unusually fragmented biodiversity hotspots of the Oriental Region have led to high levels of species diversity and endemism^46–50^. Unfortunately, the primary focus on geographical isolation, rather than on causes and consequences of adaptation and divergence in this biogeographically complex landscape, has historically misled taxonomists and evolutionary biologists in understanding and defining species, especially in polytypic taxa^5,52–55^. A unified approach to understanding species, and application of genomic data and detailed phylogenetic analyses, are now leading to improved understanding of species diversity and speciation patterns in relation to the long and complex biogeographic history of the fragmented land/seascape of the Oriental Region. Our study should be viewed in this light, where we were able to leverage a large genomic dataset, multiple lines of evidence using phylogenetic methods and population genetic tests, and data on mate choice and reproductive success. With these approaches, we were able to characterise taxonomic diversity and potential role of selection and local adaptation in the genomic diversification in this clade. Our analyses reinforced the notion that *P. polytes* as traditionally classified is not a single super-widespread species but a complex of three monophyletic species that exhibit hallmarks of being distinct species. These hallmarks include not only a strong phylogenetic and population genetic structure (Fig. 2, Fig. S1–2) but also considerable genome-level differentiation under selection (Fig. 4, Fig. S3), and pre- and postzygotic barriers to hybridisation such as strong assortative mate preference and low success of interspecific crosses (Fig. 3).

Our phylogenetic analysis using genome-wide SNPs produced a different species tree topology (Fig. 2A–C) compared with a topology shown recently^43^. This is because the Zhang et al. study^43^ did not adequately sample *P. polytes* (as delineated in our study), treating *P. javanus* as part of *P. polytes* instead, and including only a single sample of the true *P. polytes*. The multiple lines of evidence based on broad sampling in the range of *P. polytes* from which we demonstrated that *P. polytes* and *P. javanus* are indeed different species should help resolve this discrepancy and stabilise the taxonomic treatment in this species group. This should also be a lesson that polytypic taxa in the biogeographically complex Indo-Australian Region need to be reinvestigated carefully with respect to their systematic and taxonomic status.

It is notable that we did not find evidence in these young species for Haldane’s Rule—the observation that reproductive incompatibilities among different populations/species disproportionally hamper the production of individuals of the heterogametic sex in hybrid progeny^56,57^. Due to the low success of hybridisation and poor performance of hybrid progeny (Tables S5–S6), we did not have enough samples to effectively determine the sex ratio of hybrid offspring. It is possible that either these species have become sufficiently reproductively incompatible not to show effects of Haldane’s Rule or, more likely, that Haldane’s Rule needs to be specifically tested with respect to sex and species with further crossing experiments in this species group.

With respect to speciation, when should we expect few local genomic islands versus several widespread hotspots of adaptation and adaptive divergence? Theory predicts that a few genomic islands of adaptive divergence would be responsible for reproductive isolation among populations that are diverging in sympatry^8,58^. For speciation to occur, this divergence is expected to be rapid. Otherwise ongoing genetic exchange would prevent speciation, unless the fitness consequences of ecological selection on co-adapted traits/gene complexes are strong. Yet, well studied cases of sympatric speciation reveal genome-wide divergence even in young species pairs^59,60^, countering the idea of a very small number of genomic islands of differentiation and speciation.

In allopatric populations, a neutral theory would predict divergence across multiple unlinked traits and loci, but the tempo of this genomic divergence leading up to reproductive isolation and speciation to be rather slow, without producing hotspots of divergence under selection. However, we above showed genome-wide differentiation in allopatric populations, this differentiation showing unexpectedly widespread signatures of strong selection producing selective sweeps in hundreds of genes across the genome, in timespans as short as 1.27 my. Since genome-level divergence was not correlated with phylogenetic distance, and especially since so many genes with ecological and sexual functions showed signatures of selective sweeps in each species, it is clear that selection has played some prominent role in divergence among these allopatric species. Based on the signatures of selection and the functional space of genes under selection, it is possible that these species diverged perhaps for local adaptation such as mimicry and host use-related metabolism, and mate choice based on olfaction, wing patterns and courtship. We have not yet tested whether these genes are directly responsible for reproductive incompatibility and local adaptation for traits mentioned above, or they are in linkage with other loci involved in these adaptations. Similarly, we have not yet tested the relative strengths and signatures of genetic drift versus selection in the genomes of these species. Nonetheless, these findings offer many new hypotheses for future work on adaptive diversification and speciation in this iconic species group. Moreover, they suggest that the degree of selection for local adaptation and underlying genome-level divergence during both sympatric and allopatric speciation may have been underestimated. They also suggest that both sympatric and allopatric speciation events may result in widespread hotspots of genomic divergence, and not just divergence in a few genomic islands, contrary to the common expectation for allopatric speciation.

Although hundreds of recent studies have looked at the genetic basis of adaptation and divergence that pull populations and species apart, genome-level characterisation of adaptive differentiation in sister species is rarely attempted. As a result, there are few benchmarks available to compare the tempo and extent of genomic divergence under drift versus selection at advanced stages of isolation and divergence. Taking advantage of the natural experiment of diversification in these allopatric species, our work offers such a benchmark for future studies. How factors such as speciation in sympatry, secondary contact and reinforcement, and adaptive introgression, change the genome-wide signatures of selection and divergence are just beginning to be explored^59–64^. Our study offers a contrast in allopatric divergence, and might serve as a further impetus for investigations in this direction. This work should also further make this species group attractive for systematic, evolutionary and developmental genetic studies in the future.

## Acknowledgements

We thank Deepa Agashe, Chris Wheat, Jim Mallet, Ilik Saccheri and Claudius Kratochwil, for comments on the project and manuscript; Tarun Karmakar, Anuj Jain, Sarika Baidya, Arjan Basu Roy, Adam M. Cotton, and Hariprasath for contributing samples; NCBS Sequencing Facility, AgriGenome, and Genotypic for whole-genome sequencing; and Biodiversity Lab Research Collections for sample storage facilities. Wild-caught samples were obtained largely from the NCBS-TIFR campus/field stations and private lands, and from wildlife sanctuaries and national parks in India under the following research and collection permits issued by the state forest departments in Karnataka (permit no. 227/2014-2015 dated 2015/04/16), West Bengal (permit no. 2115(9)/WL/4K-1/13/BL41, dated 2013/11/06; and permit no. 1107/42/2W-705/18, dated 2018/05/07), and Meghalaya (permit no. FWC/G/173/Pt-II/474-83, dated 2014/05/27), for which we thank the Principal Chief Conservator of Forest, Deputy Conservators of Forest, Wildlife Wardens and field officers of those states.

## Author contributions

RD and SB generated and analysed genome and genotype sequence datasets; RD and MK performed selection analyses; RD analysed mate choice and reproductive success data; AG and SB performed phylogenetic analyses; AG performed species delimitation analyses; KK conceived and directed the project, designed research, performed experiments for mate choice and reproductive success; and wrote the paper with contributions from RD. All authors approved the final manuscript version.

## Additional Information

### Supplementary Information

is available for this paper.

### Data Availability Statement

Raw data for mating experiments are given in Tables S2–S4. All the sequence data used to prepare figures and tables will be submitted to NCBI, and phylogenetic trees will be submitted to Dryad, after the manuscript is initially accepted. Accession numbers will be provided in the final manuscript after initial acceptance.

## Methods

### Sample collection, sequencing and SNP calling

We performed whole genome sequencing of wild-caught samples preserved in 100% ethanol, of four species in the *polytes* group (*Papilio polytes, P. alphenor, P. javanus* and *P. protenor*), supplementing these with previously published genome sequences^43^ from the SRA database for our analysis (sample details in Table S1). We obtained *P. alphenor* and *P. javanus* from native commercial butterfly breeding facilities in the Philippines and Java, respectively. We extracted DNA from thoracic muscles of each butterfly using the QIAGEN DNeasy blood and tissue kit. We quantified extracted DNA using Qubit fluorometric quantification and prepared libraries using Illumina TruSeq DNA PCR-free library preparation kit. We sequenced them on Illumina HiSeq 2500 using 2×100 PE runs. We checked quality of sequences and downloaded genomes using FastQC, aligned them to the reference *P. polytes* reference genome from NCBI (Ppol_1.0, genome version 1.0, annotation version 1.0)^45^ using the BWA aligner^65^. We marked and removed duplicates from aligned files, merged files from the same sample, performed indel realignment, and called SNPs with haplotype caller in the GATK pipeline^66^. We filtered the resulting SNPs with the recommended parameters, retained samples with an average coverage of 9X across the genome, and obtained a set of 27,965,662 SNPs for the *polytes* species group. We filtered this SNP dataset to remove monoallelic sites and sampled SNPs every 5,000 bases to obtain a set of 30,760 SNPs that could be used for phylogenetic analysis with moderate run times.

### Phylogenetic reconstruction and species delimitation

We reconstructed Bayesian phylogenies with MrBayes 3.2.7a^67^ using four datasets: (1) 30,760 genome-wide SNPs, (2) 17 standard phylogenetic markers including both nuclear (*thiolase, carbamoyl-phosphate synthetase 2, aspartate transcarbamylase, and dihydroorotase (CAD), catalase (CAT), dopa decarboxylase (DDC), glyceraldehyde 3-phosphate dehydrogenase (GAPDH), isocitrate dehydrogenase (IDH), malate dehydrogenase (MDH), ribosomal protein S2 (RPS2), ribosomal protein S5 (RPS5), hairy cell leukemia protein 1 (HCL), elongation factor-I alpha (EF1-a)*, and *wingless*) and mitochondrial genes (*cytochrome c oxidase-I, tRNA leucine, cytochrome c oxidase II, 16S* and *ND5*)^68^, (3) a subset of the 17 markers including only the 12 nuclear genes, (4) a subset of the 17 markers including only the five mitochondrial genes. We extracted DNA sequence data for datasets 2–4 from whole genome sequences of 29 samples using bcftools-1.6 mpileup^69^: three samples each for the six species in *polytes* species group. We compiled the genome-wide SNP dataset from 48 samples: 37 samples from the *polytes* species group—five samples of *P. alphenor*, eight samples of *P. ambrax*, four samples of *P. phestus*, five samples of *P. javanus*, nine samples of *P. polytes* and six samples of *P. protenor*—and 11 outgroup species. We aligned DNA sequence data in datasets 2–4 using the MUSCLE algorithm on MEGA X^70^ and concatenated them to get a final data block of 17,450 bp for dataset 2, 12,087 bp for dataset 3, and 5,363 bp for dataset 4. Since most of these markers are housekeeping genes and therefore well-represented, we had adequate coverage for the SNPs called. We used PartitionFinder 2.1.1^71^ to choose the best partitioning scheme and models of evolution under Bayesian information criterion (BIC) for all four datasets. PartitionFinder suggested a single partition for dataset 1 (GTR), 12 Partitions for dataset 2 (GTR+I, HKY+I, HKY+G, K80, HKY+G, K80+I, GTR+I, HKY+I+G, GTR+I+G, HKY+I+G, GTR+G and HKY+I+G), six partitions for dataset 3 (GTR+I, HKY+I, HKY+G, K80, HKY+G and K80+I), and six partitions for dataset 4 (GTR+I+G, HKY+I for 2, HKY+G, HKY+I, GTR+G and HKY+I+G) (Dryad DOI for tree files will be submitted in the final submission after the initial acceptance of the manuscript). We ran the MrBayes analysis for one million generations for dataset 1 and two million generations for datasets 2–4. All four datasets had two independent runs with four chains each with sampling done every 1,000 generations. We used a split frequency below 0.01 and visual inspection of the run parameters on Tracer v1.7.1^72^ to ensure stationarity. We built consensus trees after discarding 25 percent of the samples as burnin. We used FigTree v1.4.3^73^ to view and edit all the phylogenies.

We used coalescent-based SNAPP (SNP and AFLP Package for Phylogenetic analysis) to compare alternative species models^74^, and mPTP (multi-rate Poisson Tree Processes) to determine the most supported species partition scheme by modeling branching events based on number of mutations^75^. We performed SNAPP analysis with SNAPP v 1.5.0^74^ on Beast2 v 2.6.2^76^ using the dataset of 30,760 genome-wide SNPs. We used custom scripts in R to convert SNP data to the SNAPP binary format. Due to computational limitations, we used a reduced dataset of three randomly selected samples per species to reconstruct a phylogeny using whole-genome SNPs. We tested six alternative species models (hypotheses) as shown in Table 1. For every species model, we set default model parameters: a gamma distribution with *alpha* = 2 and *beta* = 200 for the speciation rate prior of the Yule model *(lambda)* and a gamma distribution with *alpha* = 1 and *beta* = 150 for the rate prior using Beauti V 2.6.2^76^. We ran each model for 20 million generations, sampling every 2,000 generations. For the path sampling analysis, we used 48 steps, an alpha of 0.3 and a burnin of 0.25. We calculated Bayes Factor values from the Marginal Likelihood Estimates (MLE) produced by the path sampler to compare each model against the null hypothesis that *P. polytes, P. javanus* and *P. alphenor* are different species (model A). We carried out both ML and MCMC analysis on mPTP (v.0.2.4) using the multi-rate option. The MCMC analysis was run for 100 million generations, sampling every 10,000 generations in 10 independent runs. The first 2 million samples were discarded as burnin. We performed both the ML start and random start options. We examined convergence by assessing the plot of log-likelihood against MCMC iteration.

From the tree topology recovered (*protenor*,((*alphenor*,(*phestus, ambrax*)),(*polytes, javanus*))) (Fig. 2A–C), we additionally estimated time of divergence between the species splits based on the time calibrated phylogeny. The estimated node ages, along with their 95% highest posterior density (HPD), were:

> *protenor* vs other *polytes* group species: 10.2 my (HPD: 11.88 to 8.36 my)
>
> (*alphenor*,(*phestus, ambrax*)) vs (*polytes, javanus*): 4.69 my (HPD: 5.76 to 3.64 my)
>
> *alphenor* vs (*phestus, ambrax*): 3.56 my (HPD: 4.53 to 2.62 my)
>
> *phestus* vs *ambrax*: 1.95 my (HPD: 2.74 to 1.26 my)
>
> *polytes* vs *javanus*: 1.27 mya (HPD: 1.83 to 0.75 my).

### Genetic divergence between the species of *polytes* group

To find genome-wide genetic divergence between *polytes* group species, we grouped the 27 million genome-wide SNPs by species and calculated Weir-Cockerham’s Fst (in 10 kb windows) for pairwise comparisons between *P. polytes, P. javanus, P. alphenor, P. phestus* and *P. ambrax* using vcftools 0.1.13^77^. We used ADMIXTURE to estimate ancestry and population structure in the *polytes* group species using biallelic sites from the 27 million SNPs^78^. We tested ADMIXTURE runs with ancestral populations (K) ranging from 2 to 13. We plot here results of three runs with the lowest cross-validation error (K=5, K=6 and K=7; Fig. S2).

### Mate choice and hybridisation success experiments

We maintained pure-breeding populations of *polytes, javanus* and *alphenor* in separate cages. We allowed males and females from each population to mate freely and raised the progeny in the common population-specific cages. We monitored a subset of naturally occurring intra-population matings to study mating success within the population.

We set up an experiment to study inter-population mate choice and mating success in a separate cage, in which individually marked unmated males and females from each population were allowed to choose between potential mates from the three populations. These males and females were maintained in this cage from eclosion, i.e., they had no prior experience of individuals of their own population over individuals from other populations. We thus excluded the possibility of early exposure to olfactory and other signals of any particular population before they could choose between potential mates, which could otherwise bias their subsequent mate choice. This ensured that individuals were choosing mates based purely on instinctive mate preference, and not early exposure. We set up this mixed population with an equal number of males and females from each pure-breeding population.

Separately, we set up experimental hand-paired matings, in which we individually selected 3 to 5-day old unmated males and females. We paired them by gently squeezing the abdominal end and opening up claspers of a male and aligning its genital opening with that of the female, and slowly releasing the male while his claspers closed on to the female genital opening. Successful hand-pairings remove the opportunity of butterflies to court and behaviourally choose mates, and only measure the post-mating success of a pair. Such hand-pairings are possible in *Papilio*, they are often successful when done by experienced people, and have been used in hybridisation experiments in the past^79^. We noted down the duration of mating, the number of eggs laid, and stages of metamorphosis until which the progeny of the hand-paired butterflies developed successfully. All statistical tests were performed using R. We set up hand-pairings of individuals from within and across populations to compare the success of hand-paired matings. We maintained each mated female in a separate small cage in which it was allowed to lay eggs, and we raised its brood separately.

We did not include in the analysis short-lived butterflies, i.e., those that died before reaching maturity to mate (usually within the 1–3 days of eclosion) or those females that died before being able lay eggs (usually within a day after mating). Because of the low numbers, we did not attempt to study species x sex effects on the success of hybrid matings, or mating and breeding success of hybrid progeny.

### Estimation of signatures of selection in the *P. polytes* species group

To identify loci showing signatures of selection in the genomic dataset, we used a composite measure of selection (de-correlated composite of multiple signals, i.e., DCMS)^80,81^ and scanned the genomes for selective sweeps. We used the 27 million SNP dataset and calculated mu statistic using Raised Accuracy in Sweep Detection (RAiSD, v.2.5^82^) to identify loci that have experienced selective sweeps. We calculated Tajima’s D, nucleotide diversity and Fst using vcftools 0.1.13^77^ and H1 and H12 using published scripts (https://github.com/ngarud/SelectionHapStats)^83^ for the *polytes* group species and estimated a composite metric DCMS^81^ to find SNPs showing signatures of selection. We used a conservative cutoff and considered SNPs in the 99.5 percentile for both our methods and annotated them. We obtained a large number of hits with RAiSD compared to DCMS, which is expected since RAiSD is known to have a high rate of false positives depending on background selection^82^, with no overlapping hits detected by both the methods. These SNPs lay within annotated genetic elements as well as intergenic regions that lack annotation features. Genes with functional significance potentially associated with local adaptation in each species are summarised in Table S10. We mapped the genes under selection in each species and the genes common across the five mimetic species to the *Bombyx mori* genome^84^ using BLAST+ to identify their chromosomal locations. We used Circos-0.69-9^85^ to map these genes across chromosomes for each species, assuming synteny between *P. polytes* and *Bombyx mori* genomes.

We were unable to annotate with reasonable confidence genomic elements such as enhancers and other intergenic regions that may play significant roles in ecological and local adaptation, since most of them remain unidentified in these butterflies. A large number of loci that showed signatures of intense selection fell in these intergenic regions. This factor is responsible for the discrepancy in the number of total positions under selection versus the number of such loci in annotated genes (Fig. 4C).

### Characterising the functional space of genes under selection

We cross-referenced gene function for all the genes identified in our analyses to be under selection and showing the signatures of selective sweeps with the UniProt database^86^. This database provides manually annotated and peer-reviewed information on protein sequence and gene function from all organisms, although not all individual gene functions have been verified in butterflies. It is possible that some of these genes have additional functions in local and morphological adaptation in butterflies, as recently discovered for the role of *doublesex, WntA* and other genes in polymorphic mimicry and wing patterning^44,45,87,88^.

## SUPPLEMENTARY TABLES AND FIGURES

**Table S1: Sample details of wild-caught individuals used in this study**. See the Excel sheet, ‘TableS1_SampleDetails_PapilioGroup_2020-07-29.xlsx’.

**Table S2: Data for the duration and success of naturally occurring intra-population matings in pure-breeding cages**. See the Excel sheet, ‘TableS2_IntraPopulationMatings.xlsx’.

**Table S3: Data for the naturally occurring intra- and inter-population matings in a mixed cage with males and females of all three populations**. See the Excel sheet, ‘TableS3_MixedPopulationMatings.xlsx’.

**Table S4: Data for the success of hand-paired matings within and across populations**. See the Excel sheet, ‘TableS4_HandPairedMatings.xlsx’.

**Table S5:** Summaries of mate preference and reproductive success.

|  | No. of matings | Avg. duration of matings (mins) | Avg. no. of eggs laid | Avg. hatching success (%) | Avg. eclosion success (%) |
| --- | --- | --- | --- | --- | --- |
| *Papilio polytes* | 33 (84%) | 54.44±10.2 | 80.34±40.94 | 94.66±8.31 | 78.69±10.7 |
| *Papilio javanus* | 26 (93%) | 55.12±10.84 | 74.19±24.42 | 96.89±3.99 | 87.93±9.0 |
| *Papilio alphenor* | 43 (88%) | 53.63±12.8 | 91.54±46.48 | 94.49±5.93 | 79.27±8.64 |

|  | *polytes* | *javanus* | *alphenor* |
| --- | --- | --- | --- |
| *polytes* | 20 (30%) | 4 (6%) | 0 (0%) |
| *javanus* |  | 19 (29%) | 0 (0%) |
| *alphenor* |  |  | 23 (35%) |

|  | *polytes* | *javanus* | *alphenor* |
| --- | --- | --- | --- |
| *polytes* | 12; 12; 11; 11 | 12; 8; 2; 1 | 14; 11; 0; 0 |
| *javanus* | 6 | 11; 11; 9; 9 | 10; 7; 0; 0 |
| *alphenor* | 0 | 0 | 18; 18; 14; 14 |

**Table S6: Statistical results of mate preference and reproductive success**. See the Excel sheet, ‘TableS6_StatisticalTests_MatingData.xlsx’.

**Table S7: Loci experiencing selective sweep**. Annotated results of RAiSD analysis for selective sweeps. See the Excel sheet, ‘TableS6_LociUnderSelection_RAiSD_GeneAnnotations.xlsx’.

**Table S8: Loci under selection**. Annotated results of DCMS analysis for loci under selection. See the Excel sheet, ‘TableS8_LociUnderSelection_DCMS_GeneAnnotations.xlsx’.

**Table S9:** Genes under selection that show signatures of selection in more than one species. Numbers above the diagonal are for genes identified by RAiSD to be under selective sweeps, and numbers below the diagonal are for genes identified by DCMS to be under selection. There were no shared genes for three species or more in the DCMS analysis.

|  | *polytes* | *javanus* | *alphenor* | *ambrax* | *phestus* |
| --- | --- | --- | --- | --- | --- |
| *polytes* |  | 113 | 146 | 38 | 24 |
| *javanus* | 7 |  | 76 | 30 | 12 |
| *alphenor* | 2 | 1 |  | 47 | 22 |
| *ambrax* | 3 | 3 | 1 |  | 9 |
| *phestus* | 2 | 4 | 2 | 0 |  |
| No. of genes shared between *polytes*, *javanus* and *alphenor*: 43 |  |  |  |  |  |
| No. of genes shared between *polytes*, *javanus* and *phestus*: 4 |  |  |  |  |  |
| No. of genes shared between *polytes*, *javanus* and *ambrax*: 11 |  |  |  |  |  |
| No. of genes shared between *polytes*, *alphenor* and *phestus*: 8 |  |  |  |  |  |
| No. of genes shared between *polytes*, *alphenor* and *ambrax*: 16 |  |  |  |  |  |
| No. of genes shared between *javanus*, *alphenor* and *phestus*: 4 |  |  |  |  |  |
| No. of genes shared between *javanus*, *alphenor* and *ambrax*: 16 |  |  |  |  |  |
| No. of genes shared between *alphenor*, *phestus* and *ambrax*: 3 |  |  |  |  |  |
| No. of genes shared between *javanus*, *phestus* and *ambrax*: 3 |  |  |  |  |  |
| No. of genes shared between *polytes*, *phestus* and *ambrax*: 2 |  |  |  |  |  |
| No. of genes shared between *polytes*, *javanus*, *alphenor* and *phestus*: 3 |  |  |  |  |  |
| No. of genes shared between *polytes*, *javanus*, *alphenor* and *ambrax*: 7 |  |  |  |  |  |
| No. of genes shared between *phestus*, *javanus*, *alphenor* and *ambrax*: 2 |  |  |  |  |  |
| No. of genes shared between *polytes*, *javanus*, *alphenor* and *ambrax*: 1 |  |  |  |  |  |
| No. of genes shared between *polytes*, *alphenor*, *phestus* and *ambrax*: 1 |  |  |  |  |  |

**Table S10:** Genes involved in physiological and developmental processes in genomic regions experiencing selection in the *polytes* species group. These could potentially be associated with local adaptations in each of the five species. The complete list of annotated genes showing signatures of selection is given in Tables S7–8.

| Genes unique to *Papilio alphenor* | Function |
| --- | --- |
| chromatin-remodeling complex ATPase chain Iswi | gene expression, larval blood cell development, ecdysteroid signaling, metamorphosis |
| geranylgeranyl pyrophosphate synthase | carotenoid synthesis, metabolism |
| bunched | transcription, neurogenesis, eye development, oogenesis, development |
| white | transport, eye pigmentation, ommochrome biosynthesis |
| attractin | signaling, development, pigmentation |
| bone morphogenetic protein receptor type-2-like | signaling, morphogenesis, patterning, development |
| endochitinase A-like | metabolism, response to fungal pathogens |
| inhibin beta B chain | signaling, neuronal morphogenesis, optic lobe development, growth |
| cytoplasmic FMR1-interacting protein | neurogenesis, eye development |
| turtle | motor control, neuronal activity, eye development, neurogenesis |
| opsin-2-like | UV opsin, phototransduction, potential mate recognition |
| larval cuticle protein LCP-17-like | structural component of cuticle |
| PBAN-type neuropeptides-like | pheromone synthesis, signaling |
| serine protease easter-like | embryonic development, dorsoventral patterning, proteolysis, signaling |
| SH3 domain-binding protein 5 homolog | patterning, embryonic development, morphogenesis, signaling |
| phosphatase PP2A 55 kDa regulatory subunit | signaling, patterning, cell cycle, circadian rhythm |
| cytochrome P450 6B4-like | enables feeding on furanocoumarin producing plants, detoxification of xanthotoxin |
| timeless | circadian rhythm, stress response, transcription |
| rho GTPase-activating protein 7 | signaling, vesicle dynamics, neuronal activity, eye development |
| transcription cofactor vestigial-like protein 4 | transcription, negative regulation of Wnt and Hippo signalling |
| nuclear factor NF-kappa-B p110 subunit | neuronal regulation in wing disc, immunity, apoptosis |
| aromatic-L-amino-acid-decarboxylase | cuticle pigmentation, wing development, dopamine synthesis |
| homeobox-protein-homothorax | patterning, eye and leg development |
| LIM/homeobox-protein-Lhx1 | eye and leg development, patterning |
| N-acetylgalactosaminyltransferase-6-like | splicing, sex determination |
| protein-NDRG3 | spermatogenesis, signalling, negative regulation of cell growth |
| transcriptional-coactivator-yorkie | expresses in wings, regulated by hippo signalling, organ morphogenesis |
| Genes unique to *Papilio javanus* | Function |
| adenosine 3'-phospho 5'-phosphosulfate transporter 1-like | transport, dorsoventral patterning |
| cuticle protein 8-like | cuticle |
| DE-cadherin | cell adhesion, development, morphogenesis |
| desert hedgehog protein B | signaling, Patterning in frogs, development |
| exostosin-2 | morphogen (Hh, Dpp and wg) movement, signaling |
| F-box/LRR-repeat protein 4 | circadian cycle, rhodopsin signaling |
| glucose dehydrogenase [FAD, quinone]-like | cuticular modification |
| rootletin-like | sound perception, response to stimulus |
| longitudinals lacking protein-like | embryonic development, homeotic gene regulator (Scr and Ubx), transcription, patterning |
| male-specific lethal 3 homolog | chromatin remodeling, transcription, dosage compensation |
| scarlet | transport, ommochrome biosynthesis, eye pigmentation |
| angiotensin-converting enzyme | metamorphosis, muscle activity, proteolysis |
| endocuticle structural glycoprotein ABD-5-like | cuticle |
| stromal interaction molecule homolog | wing morphogenesis, calcium transport |
| cyclic AMP response element-binding protein A-like | Transcriptional, cuticle development, patterning |
| cytochrome P450 4C1-like | metabolism of insect hormones, insecticide resistance |
| knirps-related protein-like | segmentation, transcription, embryonic development |
| orexin receptor type 2-like | Transcription, patterning, cuticle development |
| son of sevenless | development, sevenless signaling pathway, signaling |
| esterase FE4-like | insecticide resistance |
| unc-80 homolog | circadian rhythm, transport |
| GTP-binding protein 2-like | eye morphogenesis, transport |
| Wnt-4-like | Wnt signaling pathway, signaling, development |
| cyclic AMP response element-binding protein A-like | memory, circadian rhythm |
| longitudinals lacking protein | transcription, neurogenesis, gonad development, development, apoptosis, eye development |
| cAMP-specific 3',5'-cyclic phosphodiesterase | cognition, circadian rhythm |
| C-terminal-binding protein | development, segmentation, interacts with hairy |
| cyclic AMP response element-binding protein A-like | memory, circadian rhythm |
| homeobox protein cut | sensory organ development, wing shape |
| homeobox protein Nkx-2.1-like | development, circadian rhythm |
| homeotic protein female sterile-like | transcription, morphogenesis, patterning |
| nuclear factor NF-kappa-B p110 subunit | regulates neuron specific genes in wing disc, immunity, apoptosis |
| odorant receptor Or2-like | olfaction |
| probable cytochrome P450 6a23 | insecticide and hormone metabolism |
| prominin-1-A | eye development |
| protein vestigial | wing development, wing disc morphogenesis, cell cycle, DNA replication, leg morphogenesis |
| serine/threonine-protein phosphatase PP1-beta catalytic subunit | cell cycle, circadian rhythm |
| serum response factor homolog | wing venation, development |
| zinc finger protein rotund | imaginal disc development, controls wingless expression |

| **Genes unique to *Papilio polytes*** | **Function** |
| --- | --- |
| keratin | silk fibres |
| odorant receptor 85b-like | signaling, response to stimuli, behavioral responses to 2-heptanone, amyl acetate, and butyl acetate |
| mushroom body large-type Kenyon cell-specific protein 1 | transcription, memory |
| ultraspiracle homolog | signaling, metamorphosis, ecdysone receptor |
| gustavus | embryonic development, cell migration, wing disc morphogenesis, metabolism |
| heterogeneous nuclear ribonucleoprotein 87F-like | embryonic development, eye development, cell cycle, splicing, metabolism, stress response |
| probable RNA-binding protein orb2 | memory, courtship behaviour, translation |
| neuralized | neurogenesis, transcription, eye development, embryonic development, wing morphogenesis, signaling, development |
| larval cuticle protein A1A-like | cuticle |
| homeobox protein homothorax | transcription, embryonic development, morphogenesis, segmentation, patterning, eye and leg development |
| vestigial | wing development, wing disc morphogenesis, cell cycle, DNA replication, leg morphogenesis |
| homeotic protein spalt-major | embryonic development, eye development, appendage development, transcription, wing vein morphogenesis |
| homeotic protein female sterile-like | transcription, morphogenesis, patterning |
| cyclic AMP response element-binding protein B | transcription, memory, circadian rhythm |
| odorant receptor 94a-like | response to stimulus, signaling |
| homeotic protein proboscipedia | development of mouthparts and palps, transcription |
| lysine-specific histone demethylase 1A | chromatin organization, transcription, wing vein morphogenesis |
| virilizer | splicing, cell differentiation, sex differentiation, required for splicing of Sxl, dosage compensation, cell migration |
| microtubule-associated protein futsch | cytoskeletal organization, eye development, neurogenesis, apoptosis |
| homeotic protein female sterile-like | transcription, morphogenesis, patterning |
| integrin alpha-PS2-like | ECM organization, signaling, cell adhesion, wing morphogenesis, development, muscle activity |
| interference hedgehog-like | signaling, Hedgehog pathway, eye development, wing patterning |
| centaurin-gamma-1A | memory, signaling, metabolism |
| larval cuticle protein F1-like | cuticle |
| neurogenic locus protein delta-like | signaling, Notch signaling, cytoskeletal organization, morphogenesis, development, appendage development, eye development, wing patterning |
| adenomatous polyposis coli protein-like | signaling, Wg signaling, proteolysis, eye development, neurogenesis |
| aryl hydrocarbon receptor nuclear translocator-like protein 1 | transcription, circadian rhythm, transport |
| protein pangolin | segmentation, embryonic development, Wnt signaling, signaling, wing morphogenesis, transcription, development |
| whirlin | sensory perception, development |
| centrosomin | cytoskeletal organization, spermatogenesis, development, hox gene regulation |
| circadian locomotor output cycles protein kaput-like | transcription, circadian rhythm, apoptosis |
| diuretic hormone class 2 | fluid secretion, circadian rhythm, signaling |
| gloverin-like | immune response |
| LIM domain kinase 1 | cytoskeletal organization, imaginal disc morphogenesis |
| longitudinals lacking protein | transcription, neurogenesis, gonad development, development, apoptosis, eye development |
| multidrug resistance protein homolog 49-like | transport, stress response, insecticide resistance |
| nuclear receptor subfamily 2 group E member 1-like | transcription, signaling, leg morphogenesis, metamorphosis, ecdysis, neurogenesis |
| odorant receptor Or2-like | olfaction |
| PDF receptor-like | circadian rhythm, signaling, locomotor rhythm |
| polyhomeotic-proximal chromatin protein-like | transcription, development, ecdysis |
| probable nuclear hormone receptor HR3 | signaling, metamorphosis, transcription |
| protein FAM107B | sensory perception |
| protein sevenless-like | phototransduction, signaling, visual perception |
| protein split ends-like | transcription, eye development, wing vein morphogenesis, signaling, wing disc patterning, development |
| protein ovo | transcription, cuticle, pigmentation, oogenesis, sex determination, cytoskeletal organization, differentiation |
| protein tramtrack | transcription, development, eye development, tracheal development, appendage development |
| LIM/homeobox protein Awh | transcription, development, eye development, imaginal disc development |

| **Genes unique to *Papilio phestus*** | **Function** |
| --- | --- |
| N6-adenosine-methyltransferase subunit METTL14 | splicing, sex determination |
| selenium-binding protein 1 | detoxification, metabolism |
| laccase-5 | metabolism, stress response, detoxification |
| protein spaetzle | signaling, embryogenesis, immune response, dorsoventral patterning |
| insulin receptor substrate 1 | signaling, cell proliferation, cell size, development, aging, reproduction |
| nucleoporin seh1 | transport, cell division, oogenesis, cell cycle |
| sodium/hydrogen exchanger 9B1-like | transport, sperm motility |
| trithorax group protein osa-like | embryogenesis, wing development, sensory perception, eye development, development, morphogenesis |
| probable serine hydrolase | metabolism, embryogenesis, development, detoxification |
| ras-specific guanine nucleotide-releasing factor 1-like | signaling, memory, vesicle dynamics, cell proliferation |
| tyrosine-protein phosphatase Lar | signaling, eye development |
| homer protein homolog 2 | signaling, sensory perception |
| dual specificity testis-specific protein kinase 2 | cytoskeletal organization, spermatogenesis, reproduction |
| transcriptional activator cubitus interruptus | wing development, transcription, eye development, morphogenesis, development, cell cycle, signaling |
| **Genes unique to *Papilio ambrax*** | **Function** |
| deleted in malignant brain tumors 1 protein-like | proteolysis, cell differentiation, immune response, transport |
| torso-like protein | signaling, development, embryogenesis, metamorphosis, cell differentiation |
| protein croquemort-like | cell death, immune response |
| neuropathy target esterase sws | metabolism, signaling, development, olfaction, sensory perception, cell death |
| RNA-binding protein 1 | sex determination, splicing, development |
| cytochrome P450 4C1-like | metabolism of insect hormones, insecticide resistance |
| neuroglian | cell adhesion, courtship, neuronal activity, morphogenesis |
| acetylcholine receptor subunit alpha-like 1 | transport, neuronal activity, insecticide resistance, signaling |

**Fig. S1:**
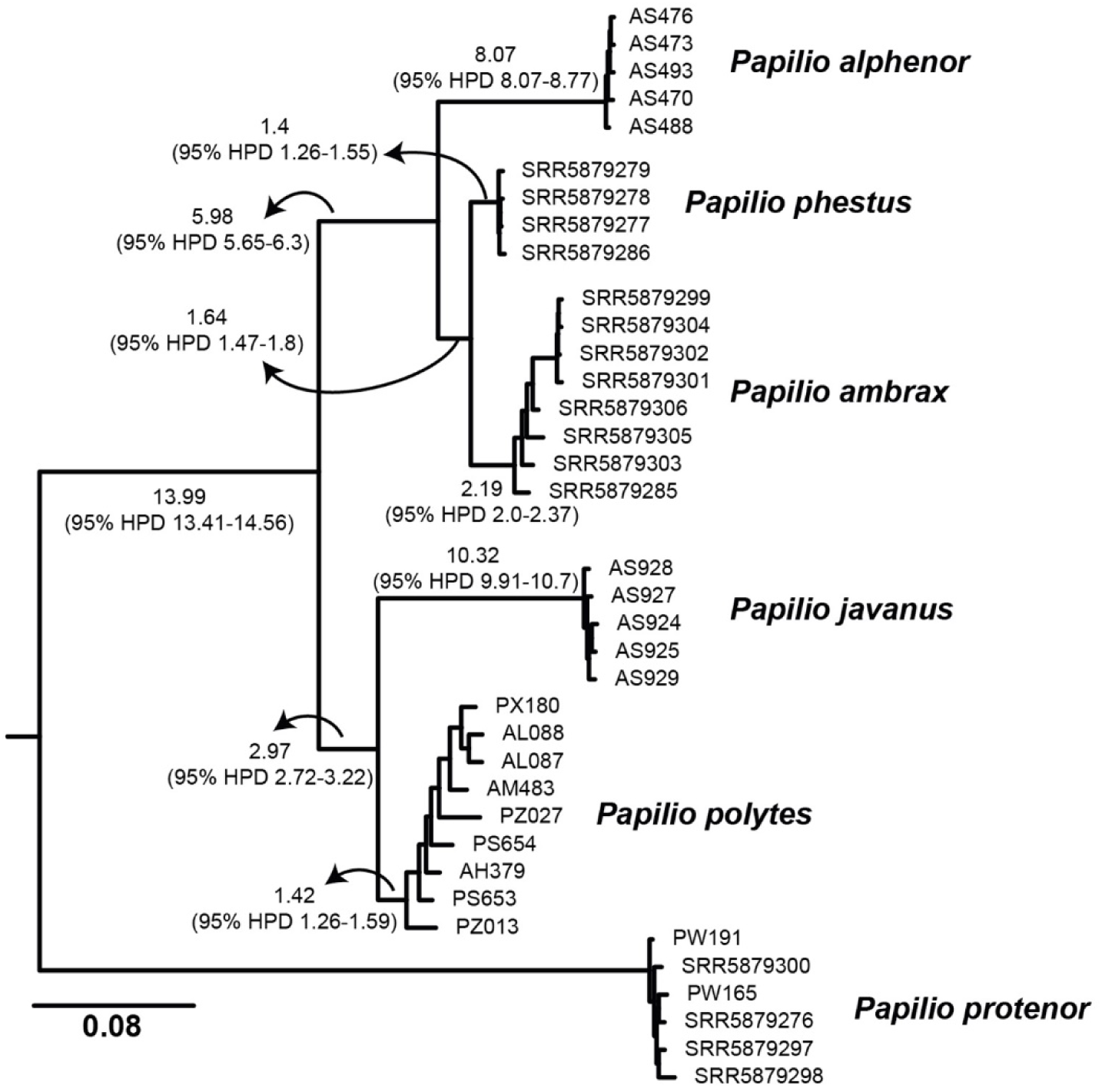
A species phylogeny of the *Papilio polytes* species group. This is an uncollapsed version of the species tree with branch statistics (percentage genetic distance), shown in Fig. 2A. Numbers with HPD are branch lengths.

**Fig. S2:**
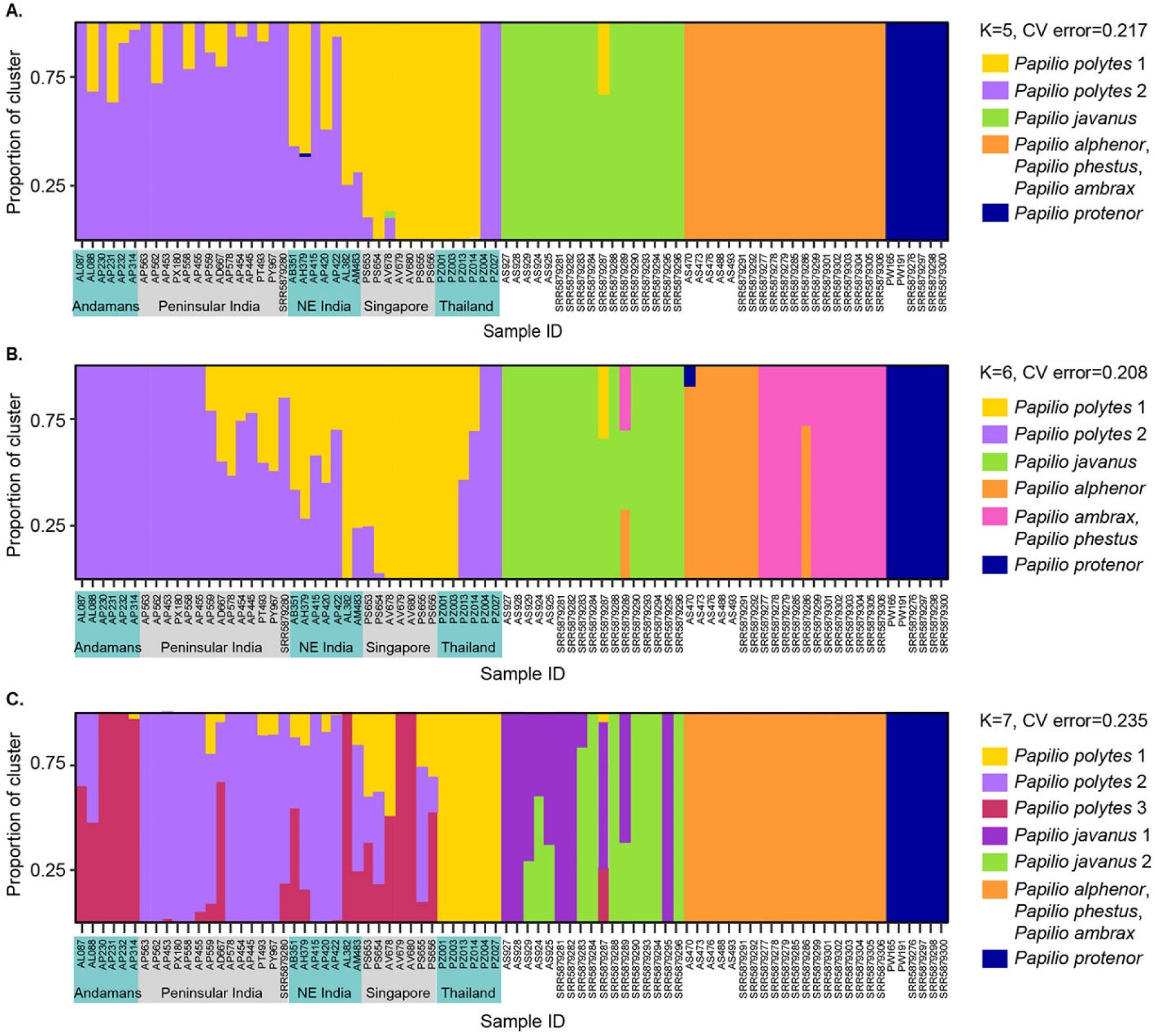
Population structure in the *polytes* species group. Different populations of *P. polytes* are represented in this ADMIXTURE analysis since this is a widespread species that occurs across the mainland and multiple islands groups. All the other ingroup species are restricted to smaller island groups. The three runs with smallest cross-validation errors are represented here (K=5,6,7).

**Fig. S3:**
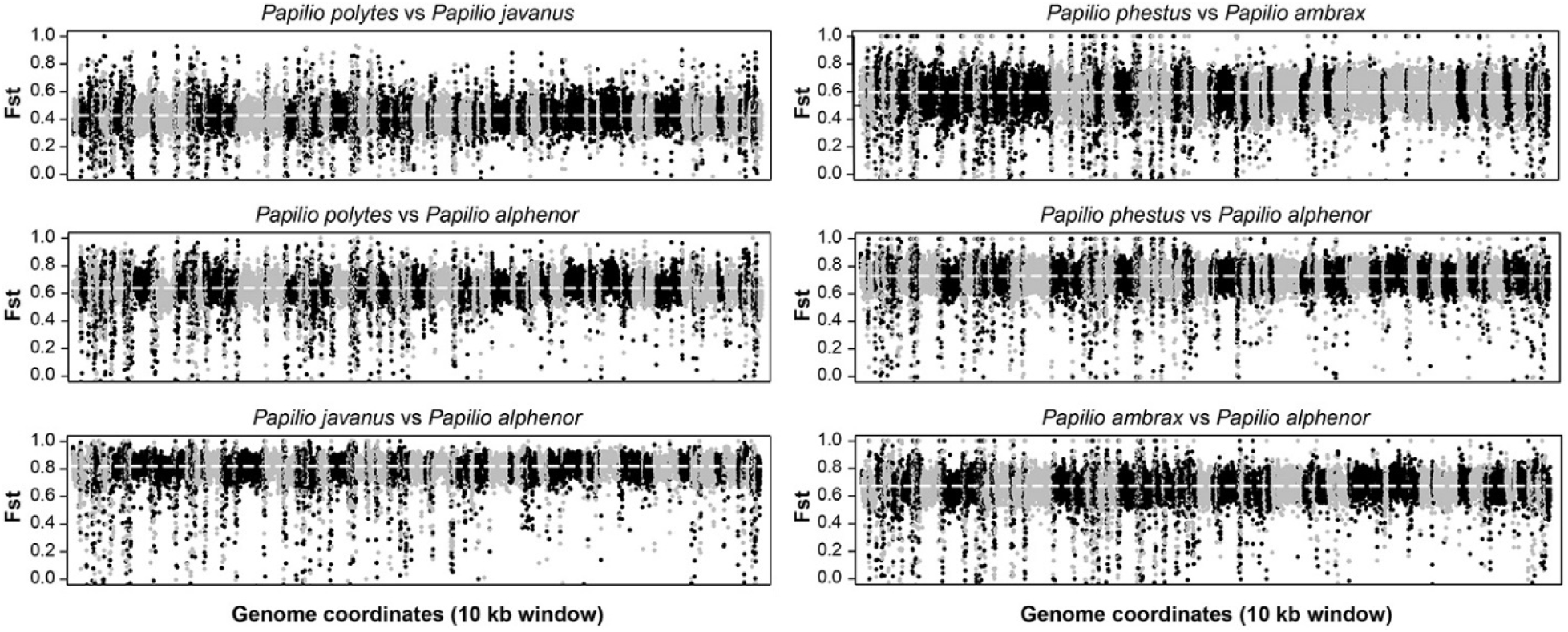
Pairwise genomic divergence between members of the *polytes* species group. Genome-wide Fst estimated in 10kb windows is shown with successive scaffolds marked black and grey, and dotted lines represent genome-wide average weighted Fst for each comparison.

